# Rational Design of the Remdesivir Binding Site in the RNA-dependent RNA Polymerase of SARS-CoV-2: Implications for Potential Resistance

**DOI:** 10.1101/2020.06.27.174896

**Authors:** Aditya K. Padhi, Rohit Shukla, Timir Tripathi

## Abstract

SARS-CoV-2 is rapidly evolving with the continuous emergence of new mutations. There is no specific antiviral therapy for COVID-19, and the use of Remdesivir for treating COVID-19 will likely continue before clinical trials are completed. Due to the lengthening pandemic and evolving nature of the virus, predicting potential residues prone to mutations is crucial for the management of Remdesivir resistance. We used a rational ligand-based interface design complemented with mutational mapping to generate a total of 100,000 mutations and provide insight into the functional outcome of mutations in the Remdesivir-binding site in nsp12. After designing 56 residues in the Remdesivir binding site of nsp12, the designs retained 96-98% sequence identity, which suggests that SARS-CoV-2 attains resistance and develops further infectivity with very few mutations in the nsp12. We also identified affinity-attenuating Remdesivir binding designs of nsp12. Several mutants acquired decreased binding affinity with Remdesivir, which suggested drug resistance. These hotspot residues had a higher probability of undergoing selective mutations in the future to develop Remdesivir and related drug-based resistance. A comparison of 21 nsp12 Remdesivir-bound designs to the 13 EIDD-2801-bound nsp12 designs suggested that EIDD-2801 would be more effective in preventing the emergence of resistant mutations and against Remdesivir-resistance strains due to the restricted mutational landscape. Combined with the availability of more genomic data, our information on mutation repertoires is critical to guide scientists to rational structure-based drug discovery. Knowledge of the potential residues prone to mutation improves our understanding and management of drug resistance and disease pathogenesis.

## INTRODUCTION

The novel coronavirus disease (COVID-19) is a highly infectious acute respiratory disease that is caused by severe acute respiratory syndrome coronavirus-2 (SARS-CoV-2). SARS-CoV-2 is closely related to other coronaviruses (CoVs), such as SARS-CoV, bat, and pangolin CoVs. However, evolutionary genomics indicates that the affinity of the Spike protein (S-protein) of SARS-CoV-2 for its human receptor angiotensin-converting enzyme 2 (ACE2) is much higher than the S-protein of other CoVs, which results in a higher infectivity rate of SARS-CoV-2 (1, 2). The genome of SARS-CoV-2 encodes four structural proteins [spike glycoprotein (S), an envelope protein (E), membrane glycoprotein (M), nucleocapsid phosphoprotein (N)] and six nonstructural proteins [open reading frame 1ab (ORF1a and ORF1ab), ORF3a, ORF6, ORF7a, ORF8, and ORF10] (3). The cleavage of ORF1a and ORF1ab polyproteins produces various nonstructural proteins (nsps) that are involved in viral transcription and replication (4). Nsp12 is the catalytic subunit and core component of the RNA-dependent RNA polymerase (RdRp) of CoVs. Nsp12 forms a complex with two additional proteins, nsp7 and nsp8, and participates in the RNA template-dependent synthesis of viral RNA in the presence of divalent metal ions (5–7). The binding of nsp12 to nsp7 and nsp8 enhances the template binding and processivity of nsp12 (5, 8). The structure of SARS-CoV-2 nsp12 shares a high homology with nsp12 of SARS-CoV, which indicates that they have similar functions and mechanisms of action (9).

The structures of the RdRp (nsp12-nsp7-nsp8) in the apo-form (8, 10) and complex form with the template primer RNA, Remdesivir, and Mg^2+^ ions were determined recently (11). The N-terminus of the nsp12 polymerase has a β-hairpin structure (residues 31-50) and an extended nidovirus RdRp-associated nucleotidyltransferase domain (NiRAN, residues 115-250) that consists of seven α-helices and three β-strands (8, 10, 12). The NiRAN domain binds at the backside of the cupped right-handed C-terminal RdRp via an interface domain (residues 251 to 365) that links the NiRAN domain to the finger subdomain of the C-terminal RdRp. The C-terminal RdRp (residues 366-920) is divided into three subdomains, finger (residues 398-581 and 621-679), palm (residues 582-627 and 688-815) and thumb subdomain (residues 816-919), which is a conserved architecture of all viral RdRps. The long finger extension of CoV RdRp intersects with the thumb subdomain to form a closed ring structure. This conformation is in contrast to the smaller loop in the RdRp of other RNA viruses, such as influenza virus, that has a relatively open conformation. Two zinc ions also bind in the conserved metal-binding motif in all CoV RdRps and play a crucial role in maintaining RdRp stability and structural integrity (8). The overall structural architecture of the apo-RdRp (without Remdesivir) and complex-RdRp (with Remdesivir) is similar, except nsp12 is in the closed conformation in apo-RdRp. The binding of nsp7 and nsp8 stabilizes the closed conformation. Remdesivir, in its monophosphate form (RMP), forms a covalent bond with the 3’ end of the RNA primer strand at the +1 position (via base-stacking interactions) and with the uridine base from the template strand (via two hydrogen bonds). RMP also interacts with side chains from Lys545 and Arg555. Two Mg^2+^ ions and one pyrophosphate are present near the bound RMP. The Mg^2+^ ions interact with the phosphodiester backbone and form part of the active site. The catalytic site of RdRP consists of seven conserved motifs (A-G) with crucial residues that are required for the activity of RdRP. The residues constituting the catalytic site and the residues involved in RNA binding are highly conserved in CoVs. Most RdRp inhibitors interact with the residues of these conserved motifs. Remdesivir is a nucleotide analog antiviral prodrug. Upon diffusion into a cell, it is converted to GS-441524 monophosphate, which is phosphorylated to the active nucleotide triphosphate form of Remdesivir (RTP). Binding of RTP leads to the inhibition of RdRp activity via nonobligate RNA chain termination (13, 14). RTP covalently binds to the +1 position and delays chain termination between positions +3 and +5. RTP inhibits the RdRp of SARS-CoV and SARS-CoV-2 with similar efficiency and mechanism of action (14). Another recent nucleoside analog inhibitor, EIDD-2801, was effective against Remdesivir-resistant SARS-CoV-2. EIDD-2801 reduced the replication and pathogenesis of CoVs in a manner similar to Remdesivir (15).

Viruses tend to adapt under drug pressure and increase the mutation rate to increase the chance per mutation of creating a beneficial mutation. This strategy ultimately leads to the development of variations for natural selection to act and create an evolving virus that is resistant to drug effects. Recent data revealed that several residues in nsp12 had mutated in SARS-CoV-2. Mutations in the Remdesivir-binding site in nsp12 may induce resistance against the drug. The present work used high-throughput computational approaches to design Remdesivir-binding sites in nsp12 and identify novel affinity-attenuating mutations. The data provide crucial insights into the residues that showed a very high tendency to undergo positive selection leading to Remdesivir resistance. The work is important to understand and manage Remdesivir resistance and may guide scientists for rational structure-based drug discovery.

## RESULTS

### Interacting residues between nsp12 and Remdesivir

The interacting residues between nsp12 and Remdesivir were obtained from the cryo-EM structure of the SARS CoV-2 RdRp. A total of 56 residues from nsp12 interacted with Remdesivir (Fig. 1A). Notably, certain nsp12 residues, such as Ala558, Gly559, Ser682, Gly683, Asp684, Ser759, Asp760, Asp761, Cys813, and Ser814, interacted with Remdesivir and template-primer RNA. A ligand-interaction diagram further revealed that the crucial residues of nsp12 were located within a 6-Å distance of Remdesivir and highlighted the metal coordination with Mg^2+^, the hydrogen bond with Asn691 and U10 of RNA, and a pi-pi stacking interaction with U20 of the template-primer RNA (Fig. 1B and 1C).

**Fig. 1.**
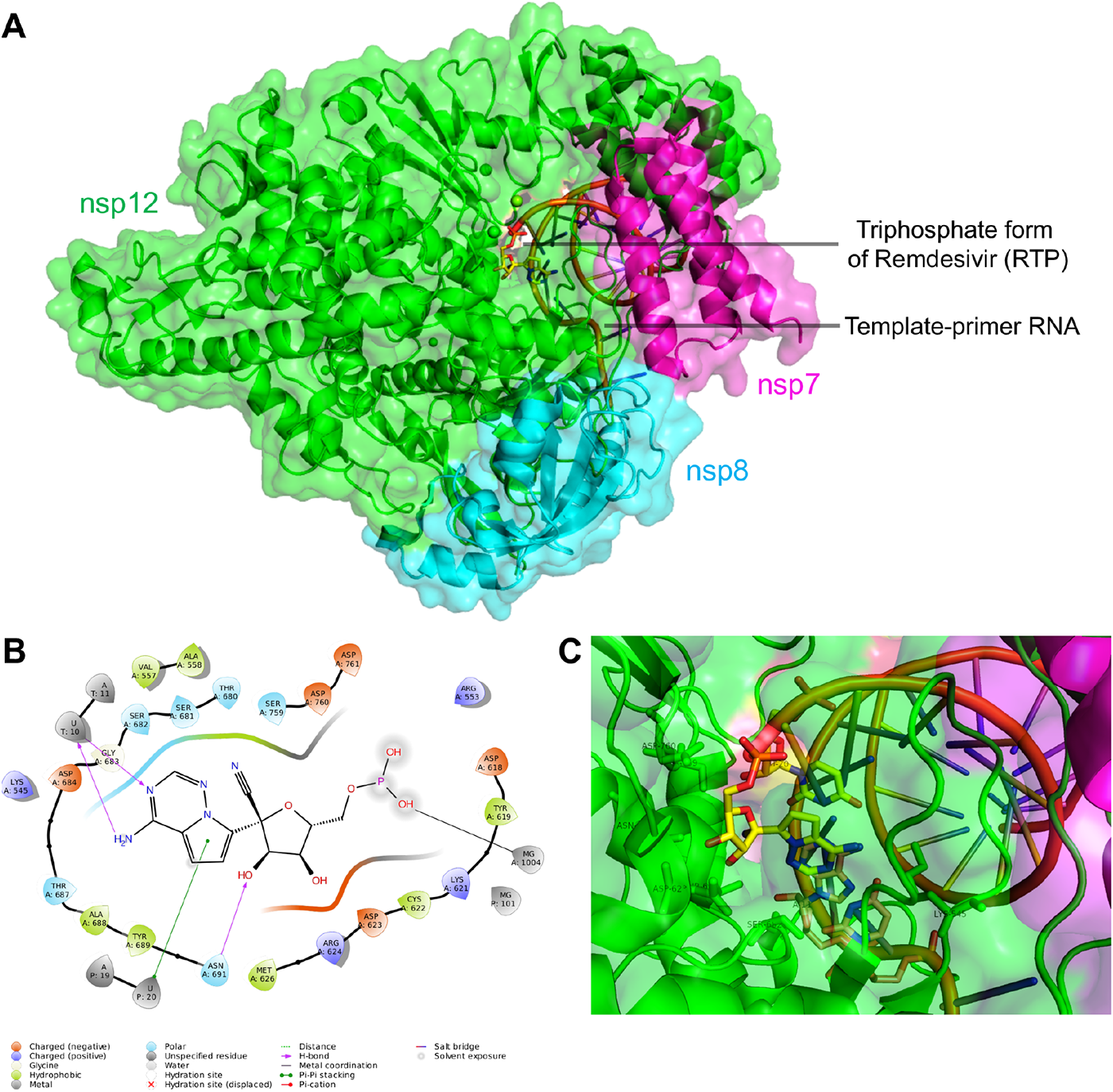
Structure of the nsp12-nsp7-nsp8 complex bound to the template-primer RNA and triphosphate form of Remdesivir (RTP), and the RTP interacting residues. (A) The cryoelectron microscopy structure of nsp12-nsp7-nsp8 complex bound to template-primer RNA and triphosphate form of Remdesivir (RTP) is shown with the catalytic Mg^2+^ ions. (B) Ligand interaction diagram of Remdesivir within a 6-Å distance of the nsp12-nsp7-nsp8 complex bound to the template-primer RNA is shown, and the types of intermolecular interactions are labeled. (C) Interacting residues of Remdesivir are labeled and shown.

### Ligand-based interface design of the Remdesivir-bound nsp12-nsp7-nsp8-RNA complex and associated physicochemical features of the complex

We performed a ligand-based interface design of the Remdesivir binding site residues of the nsp12-nsp7-nsp8-RNA complex. Remdesivir formed key contacts and interactions with 56 residues of nsp12. Notably, 10 of these 56 residues also interacted with template-primer RNA. Therefore, the 56 interface residues of Remdesivir that bound nsp12 were designed to allow the backbone flexibility of nsp12, and the residues of nsp12 other than the interface residues were repacked. A total of 50,000 designs from the interface design were computed for Rosetta’s total scores, RMSD, interface delta, and sequence identity and analyzed in detail.

First, the 50,000 designs of Remdesivir-bound nsp12 were sampled by computing the total scores versus the RMSD, and all of the designs exhibited RMSD <1.5 Å (Fig. 2A). Notably, more than half of the top-scored designs showed deviations in their RMSDs below 1 Å, which suggests that the designs did not deviate significantly from the starting complex structure during repacking, backbone movements and packing, and maintained their structural integrity even after the introduction of mutations in the 56 interface residues of nsp12 (Fig. 2A). To filter the affinity-attenuating designs from the affinity-enhancing designs, we performed a control experiment in which we only repacked (without designing) the same 56 residues of nsp12. We noted that there were designs that had poor total scores than the control and relatively higher RMSDs even compared to the affinity-enhancing designs, which suggests that the affinity-attenuating designs exhibited slight deviation in their overall structure from the starting structure and reached unfavored energetic states upon mutation.

**Fig. 2.**
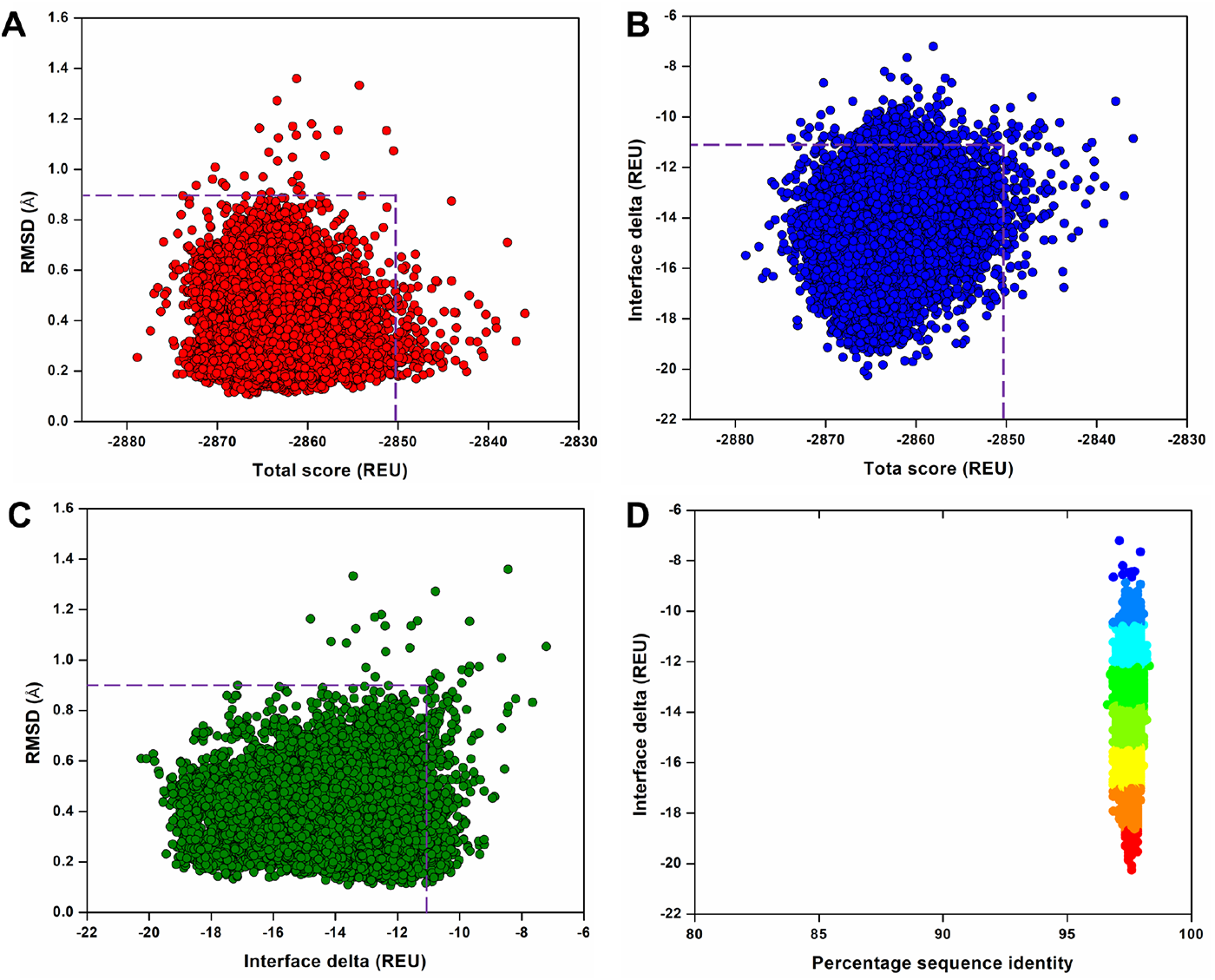
Structural and physicochemical parameters from the ligand-based Rosetta interface design experiment. (A) Rosetta total score versus RMSD of the 50,000 designs of Remdesivir-bound nsp12-nsp7-nsp8 complex bound to the template-primer RNA complex obtained from ligand-based interface design. (B) Rosetta total score versus Interface delta representing the binding affinities of the designs between Remdesivir and the nsp12-nsp7-nsp8-RNA complex. (C) Interface delta versus RMSD of the designs, where only the Remdesivir-interacting 56 residues were designed, and remaining residues were repacked. (D) Interface delta versus percentage sequence identity of the Remdesivir-interacting residues of nsp12-nsp7-nsp8-RNA complex. In panels (A-C), the dotted boxes denote the control values in which the Remdesivir-interacting residues were only repacked and not designed.

Second, the binding affinity of the designs, represented as the interface delta, was computed and plotted against the total scores (Fig. 2B). The affinity-attenuating designs had a significantly lower binding affinity than the affinity-enhancing designs, and the control in the design calculations (Fig. 2B).

Third, our computation of the RMSD versus the interface delta of the 50,000 designs revealed that the designs with lower binding affinities exhibited relatively higher RMSDs (Fig. 2C). While a significant number of the designed structures had improved binding affinities, several designed structures had reduced binding affinities with RMSDs above 1 Å as compared to the control group structures without design, suggesting that the mutations destabilized the intermolecular interactions with Remdesivir (Fig. 2C).

Finally, we evaluated the interface delta (binding affinity) of the designs to compare the affinities with the sequence identities from the native sequence. Although Remdesivir had contacts with many nsp12 residues, only a few of the nsp12 residues were more prone to mutations (Fig. 2D). The designs exhibited an overall sequence identity between 96-98%, and the lower affinity binders were less conserved than the higher affinity designs. This result suggests that although nsp12 may acquire fewer mutations at the Remdesivir binding site, certain residues are highly prone to mutations and develop resistance against Remdesivir. However, these residues can mutate whenever the virus experiences evolutionary and immune pressure during evolution.

### Validation of ligand-based interface design protocol

To validate the predictive quality and accuracy of our ligand-based interface design methodology, a previously reported Remdesivir-resistant mutant of SARS CoV, V557L, was examined (15). Val557 of nsp12 is conserved in SARS CoV-2, and it may mutate to Leu557 or any other Remdesivir-resistant mutation. To examine whether our methodology would design and rank order Leu557 as one of the low-scored designs, we scored and scanned residues in the 557^th^ position and found that V557L was ranked as a low-scored design in the design calculations (Fig. S1). This result demonstrated that our methodology was capable of scoring, rank-ordering, and filtering the resistance-enhancing mutations from that of the possible resistance-attenuating mutations.

### Sequence diversity of Remdesivir binding nsp12 designs

The 50,000 designs that were generated surrounding the Remdesivir-binding site of nsp12 were classified into affinity-attenuating and affinity-enhancing designs. The 56 interacting amino acid residues of the 100 top-scored affinity-attenuating designs were analyzed and compared with the 100 top-scored affinity-enhancing designs with a higher total score and binding affinities. A mutational landscape analysis revealed that residues Tyr456, Asn543-Ser549, Lys551, Ala554-Thr556, Gly559, Val560, Asp618-Lys621, Ala625, Gly679, Ser681, Ser682-Thr686, Tyr689, Ser692, Asn695, Ser759, Asp761, Ala762, Asn790, Glu811, Phe812, and Ser814 were sampled to almost identical residues in both groups, and they varied most at the remaining residues (Fig. 3). Notably, some residues, such as Arg553, Val557, Ala558, Trp617, Arg631, Val662, and Phe694, exhibited diverse mutations between the two groups but with a smaller number of amino acids sampled in these positions, other residues, including Ile589, Cys622, Asp623, Arg624, Met626, Thr680, Ser682, Thr687, Ala688, Ala690, Asn691, Leu758, Asp760, and Cys813, experienced more diverse sequence variations between the two groups with a relatively higher number of amino acids sampled (Fig. 3). Notably, motif A (residues 613-626) and motif B (residues 675-710) were more prone and highly susceptible to mutations, and the catalytic residues Ser759 and Asp761 (which also interact with primer RNA) were completely conserved (Fig. 3). Twenty-one residue positions of nsp12 varied between the affinity-attenuating and affinity-enhancing designs. This mutational landscape data between the affinity-attenuating and affinity-enhancing designs showed that certain amino acids and positions in the nsp12 were highly prone to mutations and even accommodated a higher number of variations. For example, the R553S, A558S, and A690D mutations that were sampled in the affinity-attenuating designs were already identified in the SARS-CoV-2 genomes, as reported in the CoV-GLUE database, which contains replacements, insertions, and deletions and were observed in the GISAID hCoV-19 sequences sampled from the pandemic (http://cov-glue.cvr.gla.ac.uk/#/replacement), which suggests that these viruses are probably resistant to Remdesivir (Fig. 3). This sequence-specific conservation and diversity of the Remdesivir-bound nsp12 designs show the design accuracy and predictive capability of our design methodology, which suggests that these hotspot residues will likely undergo selective mutations in the near future to develop Remdesivir and related drug-based resistance for the propagation of infection and survival of SARS-CoV-2.

**Fig. 3.**
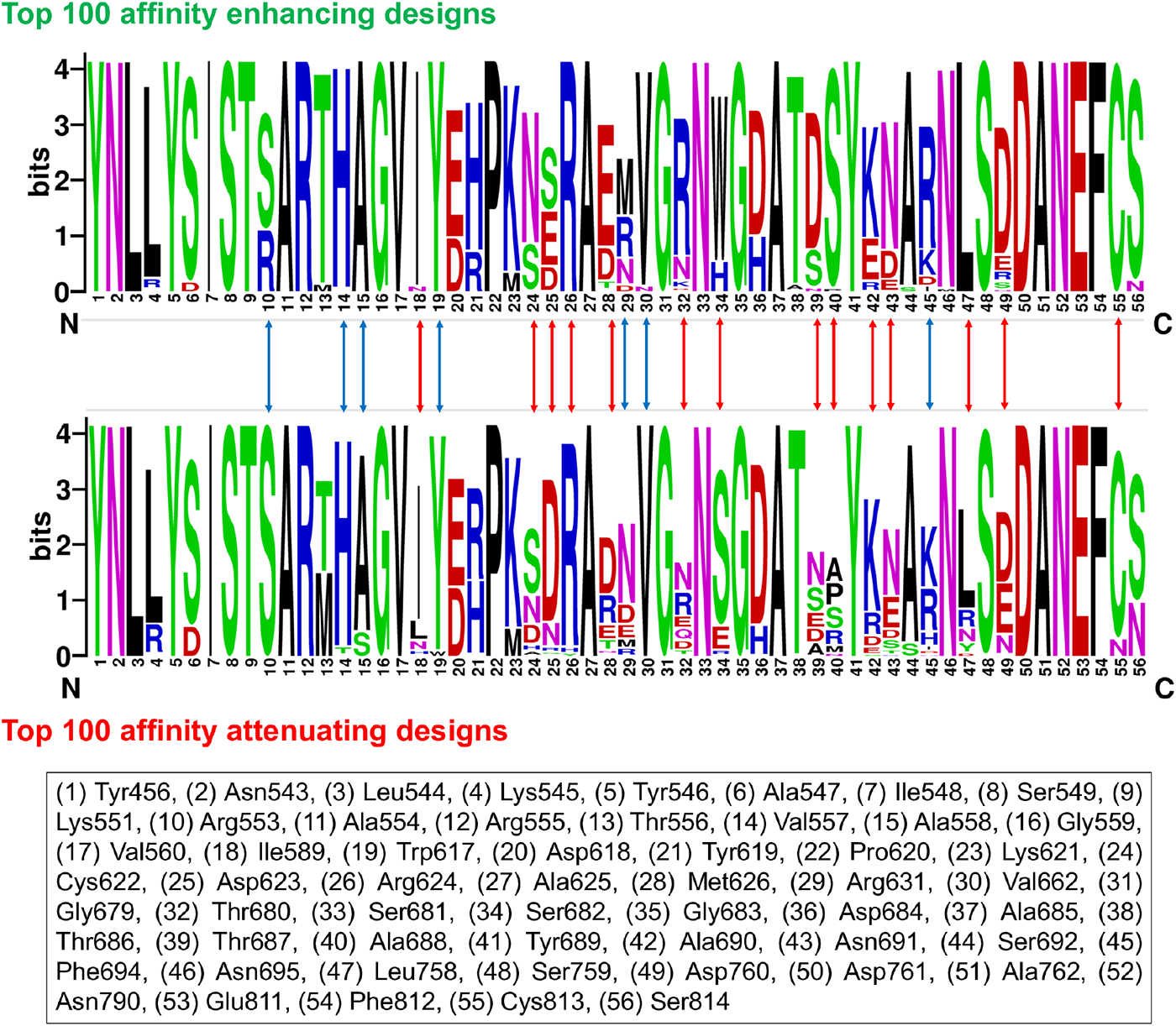
Sequence logos showing the frequency of designed Remdesivir-interacting residues. Sequence logos of the 100 top-scored affinity-attenuating versus affinity-enhancing designs generated from the ligand-based interface design of the Remdesivir-bound nsp12-nsp7-nsp8-RNA complex are shown. The integers on the X-axis represent corresponding native residues, and their identities are shown in the bottom panel. The ‘bits’ represent the overall height of the stack in the Y-axis with the sequence conservation at that position. The blue arrow shows residues that exhibited diverse mutations between the two groups but with a fewer number of amino acids sampled, and the red arrow shows residues that experienced more diverse sequence variations between the two groups with a relatively higher number of sampled amino acids.

### Binding affinity of Remdesivir-bound nsp12 designs using PRODIGY-LIG

After the Rosetta ligand-based interface design calculations, we computed the binding affinities between Remdesivir and all of the Rosetta-generated designs using PRODIGY-LIG. We found that the binding affinities of Remdesivir-bound designs ranged from −5.01 kcal/mol to −6.35 kcal/mol (Fig. S2). Several affinity-attenuating designs generated from Rosetta also had lower binding affinities in PRODIGY-LIG scoring. For the wild-type Remdesivir-bound nsp12-nsp7-nsp8-RNA complex, the binding affinity between nsp12 and Remdesivir was −6.02 kcal/mol, with a total of 2862 atomic contacts. Interestingly, while many of the top designs had a substantial increase in binding affinity, there were several affinity-attenuating designs with reduced binding affinity with Remdesivir (Fig. S2).

### Intermolecular interactions and visualization of Remdesivir-bound top-ranked affinity-enhancing and affinity-attenuating nsp12 designs

To compare how the top-scored affinity-attenuating Remdesivir-bound designs differ from the top-scored affinity-enhancing designs, we were interested in visualizing the various types of intermolecular interactions between them. The number of intermolecular interactions that determine these variations were also computed. The affinity-enhancing design formed 349 interactions, and the affinity-attenuating design formed only 260 interactions in total (Table 1). Proximal interactions, polar contacts, hydrogen bonds, and aromatic contacts were the major interactions that governed the increased or decreased affinity between Remdesivir and nsp12 (Fig. 4A and 4B). A ligand interaction diagram between Remdesivir and nsp12 for the affinity-enhancing and affinity-attenuating designs shows that the lack of a pi-pi stacking interaction between Remdesivir and His682 of nsp12 was one of the major contributors to lower binding affinity in the affinity-attenuating design (Fig. 4C and 4D).

**Fig. 4.**
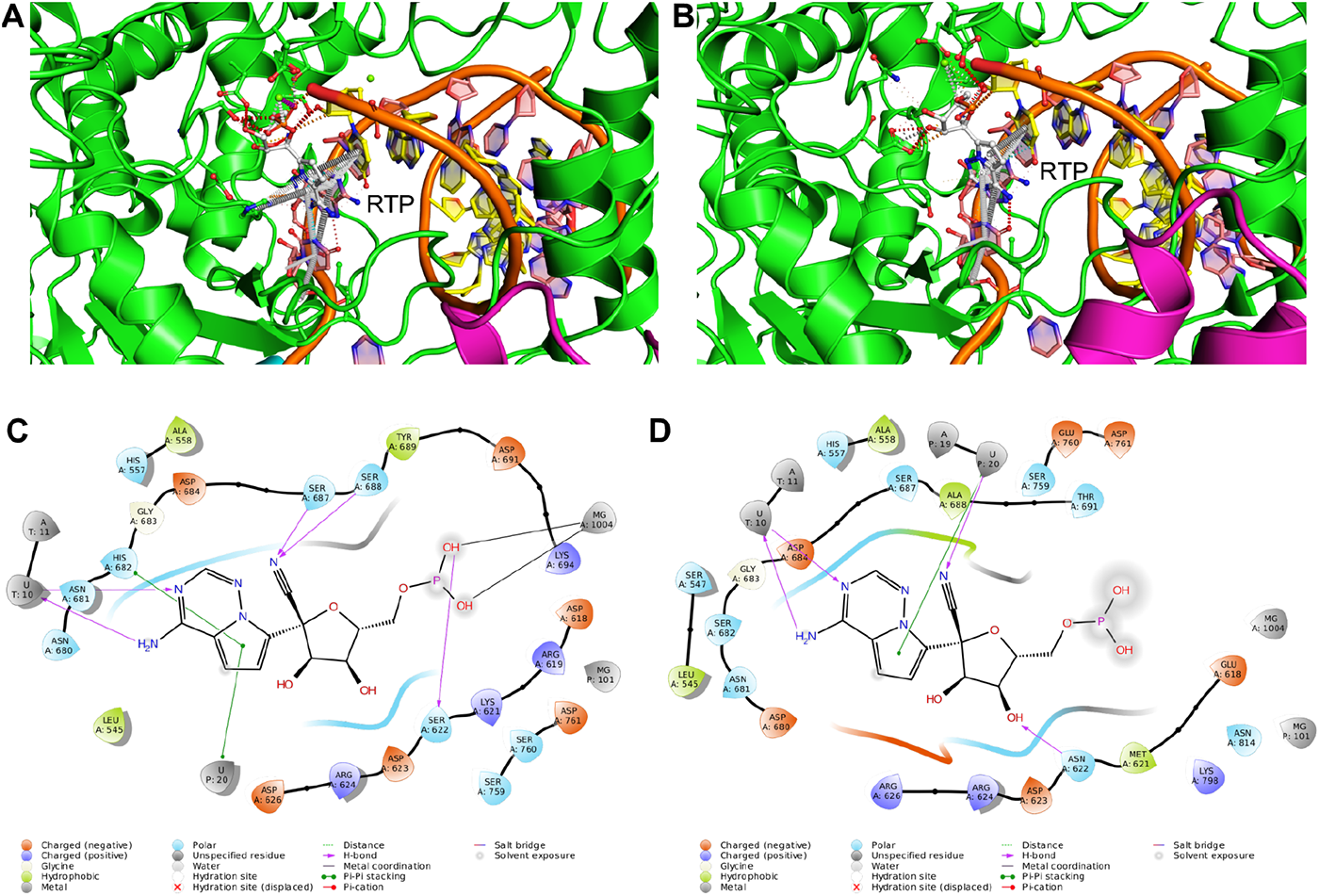
Key interactions and ligand interaction diagrams between Remdesivir and the nsp12-nsp7-nsp8-RNA complex for the top-scored affinity-enhancing and affinity-attenuating designs. Key intermolecular interactions between Remdesivir and the nsp12-nsp7-nsp8-RNA complex for the top-scored (A) affinity-enhancing designs and (B) affinity-attenuating designs are shown. Various types of intermolecular interactions, such as van der Waals interactions, proximal interactions, polar contacts, hydrogen bonds, aromatic contacts, hydrophobic contacts, carbonyl interactions, and amide-amide interactions, are shown in yellow, gray, red, white dashed, white long-dashed, green dashed, black-white dashed and blue dashed lines, respectively. 2D ligand interaction diagrams between Remdesivir and the nsp12-nsp7-nsp8-RNA complex for the top-scored (C) affinity-enhancing designs and (D) affinity-attenuating designs are shown within a 6-Å distance. Various types of intermolecular interactions are labeled as a legend.

**Table 1.** Detailed intermolecular interactions formed between Remdesivir and nsp12 binding residues and between EIDD-2801 and nsp12 binding residues in the top-scored affinity-enhancing and affinity-attenuating designs.

|  | **Remdesivir-nsp12** |  | **EIDD-2801-nsp12** |  |
| --- | --- | --- | --- | --- |
| **Types of interactions** | **Affinity enhancing design1** | **Affinity attenuating design1** | **Affinity enhancing design1** | **Affinity attenuating design1** |
| Van Der Waals interactions | 4 | 7 | 5 | 3 |
| Proximal interactions | 285 | 219 | 257 | 213 |
| Polar contacts | 17 | 10 | 12 | 8 |
| Hydrogen bonds | 8 | 3 | 8 | 2 |
| Aromatic contacts | 33 | 19 | 7 | 15 |
| Hydrophobic contacts | 2 | 2 | 5 | 4 |
| Carbonyl interactions | 0 | 0 | 2 | 0 |
| Total number of interactions | 349 | 260 | 296 | 245 |

### Comparative analysis between EIDD-2801-bound nsp12 designs to Remdesivir-bound nsp12 designs

The antiviral drug EIDD-2801 was docked into the Remdesivir binding site of SARS-CoV-2 RdRp, and EID-2801 and Remdesivir shared several common interactions with nsp12, including hydrogen bonds with Asn691 and the U10 and U20 of the primer RNA. EIDD-2801 also established a unique hydrogen bond with Ser159 (Fig. 5A and 5B). Because EIDD-2801 shared a significant number of similar interactions with Remdesivir (Fig. 5C), we performed a ligand-based interface design of EIDD-2801 binding site residues of nsp12 and found 65 contacts and interactions with nsp12. The identical design condition and methodology used in the Remdesivir-based nsp12 design was performed, and 50,000 designs from the interface designs were computed for Rosetta total score, RMSD, interface delta, and sequence identity.

**Fig. 5.**
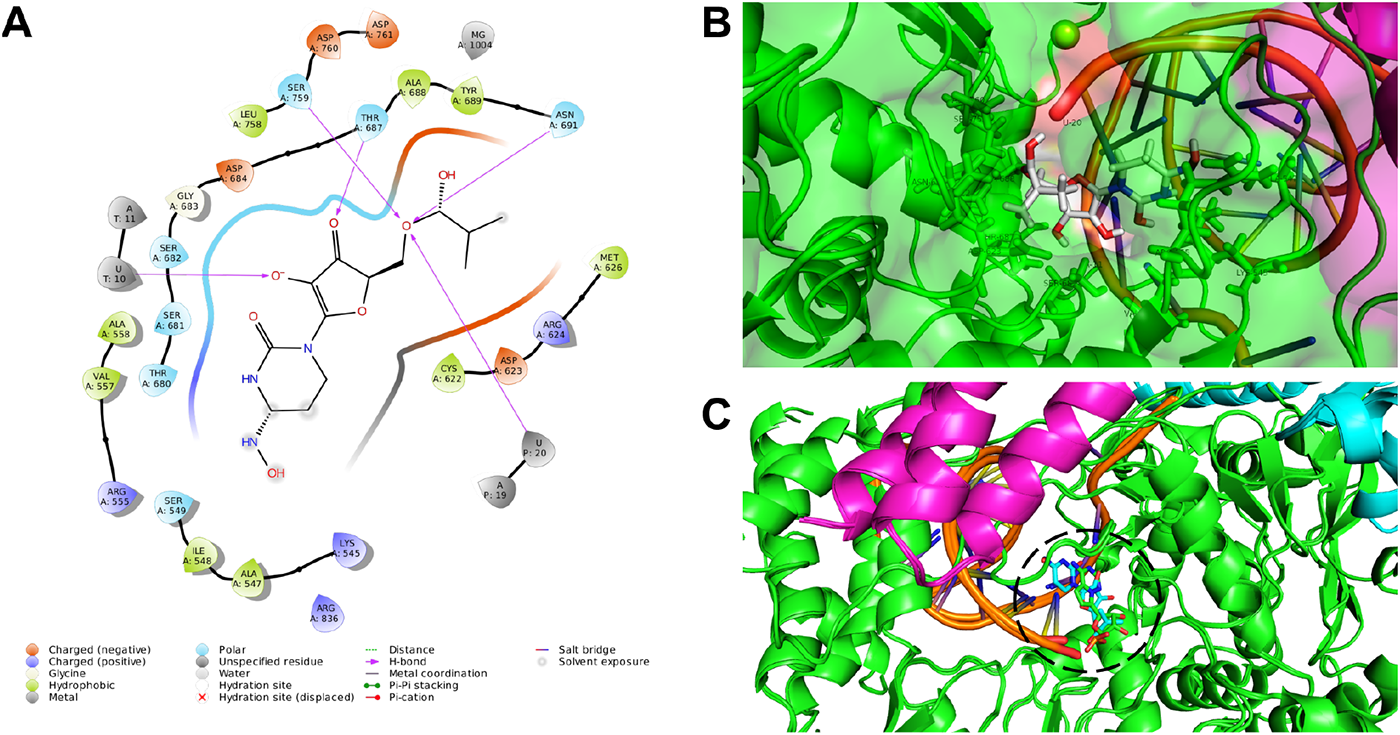
Interactions between EIDD-2801 and the nsp12-nsp7-nsp8-RNA complex and its docked pose inside nsp12-nsp7-nsp8-RNA. (A) Ligand interaction diagram of EIDD-2801 within a 6-Å distance of the nsp12-nsp7-nsp8 complex bound to the template-primer RNA is shown. The types of intermolecular interactions are labeled. (B) Interacting residues of EIDD-2801 are labeled and shown. (C) Structural superposition of the starting structure of the Remdesivir-bound nsp12-nsp7-nsp8-RNA complex and the EIDD-2801-docked nsp12-nsp7-nsp8-RNA complex is shown, where Remdesivir is shown as a green stick, and EIDD-2801 is shown as a cyan stick, and a very identical binding pose and conformation was observed.

Similar to the Remdesivir-bound nsp12 designs, we first sampled the 50,000 designs of EIDD-2801-bound nsp12 by computing the total scores versus the RMSD (Fig. S3A). We also performed a control experiment in which we only repacked the same 65 residues of nsp12 and considered the designs that were poorer in total score and binding affinity. We found that the designs with higher-energy also exhibited RMSDs far from the native (>1.0 Å). Our subsequent calculations of total score versus interface delta, which showed the binding affinity, demonstrated that the lower binding affinity designs retained relatively higher energies, and a substantial number of designs had lower binding affinity compared to the control group structures (Fig. S3B). We analyzed and plotted the RMSD versus the interface delta of the designs and observed that the designs with significantly lower binding affinities for EIDD-2801 were structurally a bit far from the native structures (Fig. S3C). More than half of the designs scored poorer than the control group, which showed the effect of mutations in the EIDD-2801 binding site of nsp12. The final evaluation of the binding affinities compared to the sequence identity of the designs revealed similar characteristics to the Remdesivir-binding nsp12 designs, in which only four residues were more prone to mutations (Fig. S3D). This result suggests that a few residues of nsp12 from SARS-CoV-2 are more susceptible to mutations at the EIDD-2801 binding site and may develop resistance via beneficial mutations to avoid evolutionary and external pressure.

The 50,000 designs that were generated surrounding the EIDD-2801 binding site of nsp12 were classified into affinity-enhancing and affinity-attenuating designs, similar to Remdesivir. The 65 interacting amino acid residues of the 100 top-scored affinity-enhancing and affinity-attenuating designs from each group were analyzed. A mutational landscape analysis revealed that residues Arg555, Val557, Val560, Tyr619, Met626, Ser682, Thr687, Ala688, Asn691, Phe694, and Asn695 exhibited diverse mutations between the two groups but with a fewer number of amino acids sampled in these positions (Fig. S4). Notably, some other residues, including Lys545 and Cys622, exhibited more diverse sequence variations between the two groups and with a relatively higher number of amino acids sampled (Fig. S4). Thirteen residue positions of nsp12 varied between the affinity-enhancing and affinity-attenuating designs compared to 21 residues in the Remdesivir-bound nsp12 designs. This mutational landscape data between the affinity-attenuating and affinity-enhancing designs shows that Remdesivir binding sites of nsp12 had a higher number of mutations, and the EIDD-2801 binding site residues of nsp12 were relatively immune to mutations.

Our binding affinity calculations using PRODIGY-LIG for the EIDD-2801-bound designs showed that the designs exhibited binding affinities that ranged from −6.26 kcal/mol to −7.27 kcal/mol (Fig. S5). With the decrease in the number of total contacts, the binding affinities were decreased. However, the difference in the binding affinity of the Remdesivir-bound designs was higher (−1.34 kcal/mol) compared to the EIDD-2801-bound designs (−1.01 kcal/mol).

We visualized the various types of intermolecular interactions between the top-scored affinity-enhancing designs of EIDD-2801 and the top-scored affinity-attenuating designs of EIDD-2801. We found that the affinity-enhancing design formed 296 interactions, but the affinity-attenuating design formed only 245 interactions in total (Table 1). Proximal interactions, polar contacts, and hydrogen bonds were the major interactions that governed the increased or decreased affinity between EIDD-2801-nsp12. A ligand interaction diagram between EIDD-2801 and nsp12 for the affinity-enhancing and affinity-attenuating designs clearly shows that the absence of a higher number of hydrogen bonds with Asn691 and Arg682 formed between nsp12 and EIDD-2801 contributed to the lower binding affinity (Fig. S6).

### Normal mode analysis of nsp12

We further computed the normal modes of nsp12 of SARS-CoV-2 RdRp to understand which portions and sites of nsp12 undergo large amplitude movements. Comparisons of these modes to our Remdesivir-binding site designs of nsp12 is crucial to understand the dynamics and flexibility of crucial nsp12 sites. First, our analysis of the displacement of each Cα atom for modes 7 to 12 revealed that residues 400-750 were mostly stable across all modes (Fig. S7). Second, our calculation of the square of the fluctuation of each Cα atom calculated from the 200 lowest nontrivial modes and normalized highlighted that nsp12 residues from 400-800 exhibited fewer fluctuations compared to other sites (Fig. 6A). Finally, our residue correlation analysis, which represented the correlated movement of the Cα atom in the nsp12 protein, showed that residues 450-780 of nsp12 undergo a highly correlated coupled motion (Fig. 6B). All of these data suggest that nsp12 residues 450-780 are highly stable and do not exhibit much fluctuation. These data correlated well with our design results, which showed that the Remdesivir-bound nsp12 designs exhibited lower RMSDs and did not undergo large conformational changes even after mutations.

**Fig. 6.**
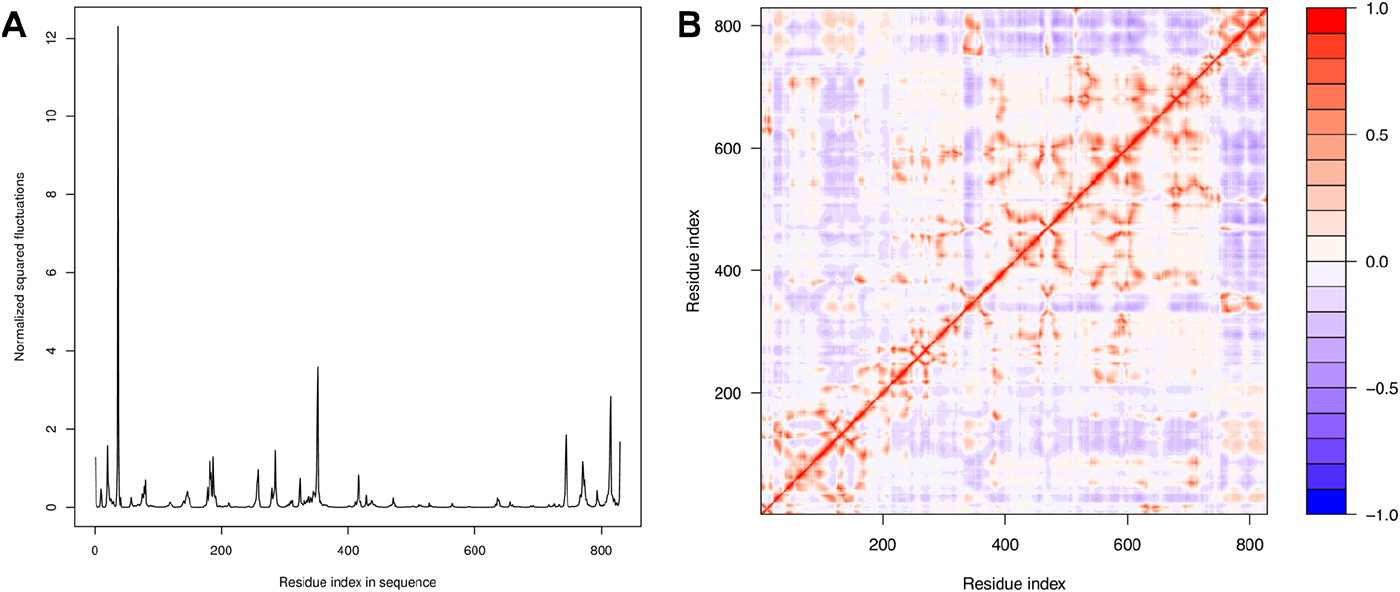
Normal mode analysis of RdRp nsp12. (A) The normalized squared fluctuations, and (B) the correlation matrix of nsp12 obtained from the normal mode analysis are shown.

## DISCUSSION

RNA viruses have a remarkable ability to adapt to new hosts, environments, and drugs via the development of genetic variations in a short time (16). The RNA viruses tended to mutate faster than DNA viruses, and single-strand viruses exhibit higher mutation rates than double-strand viruses. Mutation rates are not fixed and evolve in response to drug pressures. There are many examples of virus mutations that alter virulence or cause drug resistance. The resistance against the nucleoside analog azidothymidine (AZT) is a classic example of the importance of viral mutation. AZT was the first approved drug for human immunodeficiency virus (HIV) treatment, but HIV rapidly developed AZT-resistance mutations that led to the failure of AZT monotherapy. SARS-CoV-2 has maintained a low mutation rate, which is attributed to the 3’-to-5’ exoribonuclease activity of the nsp14 protein, which helps maintain its replication fidelity (17). The low mutation rate of SARS-CoV-2 suggests that it may evolve relatively slowly compared to other viruses, but the comparative rates of evolution also depend on how many sites are targeted by selection. However, SARS-CoV-2 often fails to infect close contacts, which suggests that there is an abundant scope to increase transmission ability in the future (18). The viral load in a patient is very high (~100,000 viral copies per mL of saliva), which suggests that every possible mutation may develop over the course of the infection (19). The occurrence of extremely high viral loads may also account for the fast-spreading nature of the COVID-19 pandemic.

RdRps catalyze the RNA template-dependent formation of phosphodiester bonds and facilitate virus replication and transcription. The nsp12 subunit of the RdRp of SARS-CoV-2 and SARS-CoV share a high homology, which suggests that its function and mechanism of action are well conserved (10, 20). RdRps are the primary targets for antiviral drug development against SARS-CoV-2, and several RdRp-targeting drugs, such as Remdesivir, Favipiravir Galidesivir, Ribavirin, and Penciclovir, are in testing stages (21–23). Remdesivir is an investigational, broad-spectrum antiviral drug that demonstrated activity against CoVs and other RNA viruses. It is being tested as a potential treatment for COVID-19 in multisite clinical trials, and it recently advanced to a phase 3 stage (24). Remdesivir is a nucleoside analog that acts as an RdRp inhibitor and targets viral replication. Few naturally occurring mutations in RdRp that led to drug-resistance were observed previously (22, 25, 26). Since its emergence in November 2019, several mutations in the RdRp of SARS-CoV-2 were reported. A total of 783 nsp12 mutations have been reported and are listed in the CoV-GLUE database, and 15 residues are in the Remdesivir-binding site. A recent study identified mutation hotspots in SARS-CoV-2 (27). They showed that viral strains with RdRp mutation had a median of 3-point mutations, and viral strains with no RdRp mutation had a median of 1 mutation. A mutation in position 14408 of SARS-CoV-2 RdRp was associated with an increased mutation rate (27). Two nsp12 mutations (F480L and V557L) conferred an up to 5.6-fold resistance to SARS-CoV (22), and these residues are conserved across all CoVs.

A detailed atomic-level understanding of RdRp interaction with Remdesivir using robust molecular technologies could revolutionize the development of new therapies to fight COVID-19 in a highly specific and efficient manner. We designed 50,000 Remdesivir-interacting nsp12 residues using high-throughput ligand-based interface designing. We also docked the oral NHC prodrug (β-D-N^4^-hydroxycytidine-5’-isopropyl ester) EIDD-2801, which is similar to Remdesivir because both drugs work by mimicking ribonucleosides, into the Remdesivir binding site of SARS-CoV-2 RdRp and designed 50,000 EIDD-2801-interacting nsp12 residues. Therefore, a total of 100,000 designs were made and analyzed in detail. The most striking result was that even after designing 56 residues in Remdesivir binding site of nsp12 and 65 residues in the EIDD-2801 binding site of nsp12, the designs retained 96-98% sequence identity, which suggests that SARS-CoV-2 attained resistance with very few mutations in nsp12. These highly mutation-prone residues/sites should be critically considered in the development of future therapeutics and during analyses of pandemic evolution. The affinity-attenuating designs showed lower Remdesivir binding, which indicates that the SARS-CoV-2 could develop Remdesivir resistance via mutations at these residues. These hotspot residues would likely undergo selective mutations in the future to establish Remdesivir and related drug resistance. Our design methodology mutated and rank-ordered specific Remdesivir-resistance mutations of SARS-CoV, which highlights its accuracy and predicting ability. The normal mode analysis and residue correlation analysis suggest that residues 400-750 are mostly stable across all modes and undergo a highly correlated coupled motion, which suggests that this region does not exhibit much fluctuation. These results correlate well with our design results, in which Remdesivir-bound nsp12 designs did not undergo large conformational changes, and retained 96-98% sequence identity even after mutations modulated the binding affinities with Remdesivir and EIDD-2801. Compared to the 21 nsp12 residues in the Remdesivir-bound designs, only 13 nsp12 residues varied in the EIDD-2801-bound designs. This result suggests that EIDD-2801 is more effective in preventing the emergence of resistance mutations due to the restricted mutational landscape and explains why EIDD-2801 is effective against Remdesivir-resistance mutations. Our cohesive and comprehensive work highlights the hotspot residues of nsp12 that have the potential to undergo mutation to develop Remdesivir resistance. Information on potential mutants could help clinicians adjust drug regimens to administer treatments earlier and more aggressively in higher-risk conditions.

We acknowledge certain limitations in the current study. Our Rosetta ligand-based interface design method (and scoring function) is set to design mutants with improved interface delta and therefore increased binding affinity to Remdesivir. However, ideally, the Remdesivir-resistance mutations are affinity-attenuating designs. To the best of our knowledge, at present, neither Rosetta nor any other programs have protocols to accurately score and predict affinity-attenuating designs to correlate with a drug resistance mutation. Nonetheless, Rosetta methodology remains the only useful design-based tool for mutational mapping and prediction of specific residues involved in enhancing or attenuating protein-ligand interaction. Therefore, to make sure that our Rosetta protocol captures the correct affinity-attenuating designs to correlate with Remdesivir-resistance mutations, we carried out control experiments by taking the wild-type and a previously known Remdesivir-resistance SARS-CoV mutation as control. We argue that the affinity-attenuating designs representing the potential hotspot residues have a higher probability of undergoing selective mutations in the future to develop Remdesivir and related drug-based resistance. Finally, owing to a lack of the structure of the RdRp-EIDD-2801 complex, the exact structural information predicted from the docked complex of RdRp-EIDD-2801 might not be precise; however, since Remdesivir and EIDD-2801 bind at the same position, the error might not be significant enough to impact our design results.

We conclude that, in the absence of detailed information on the evolutionary forces and the role of selection pressures, predicting how a virus may evolve is a challenging task, particularly during a nascent outbreak. The rapid availability of more SARS-CoV-2 genomes, most of which differ by one or few mutations, is enabling accurate investigations into spread patterns. Genomic epidemiology-based analysis is being used to control the fast-growing SARS-CoV-2 outbreak. Our prediction on the mutation repertoires complements research to understand and manage drug resistance and pathogenesis and guides development strategies to deter future cross-species transmissions.

## METHODS

### Identification of interacting residues between nsp12 and the triphosphate form of Remdesivir (RTP)

The cryo-EM structure of the RNA-dependent RNA polymerase from SARS CoV-2 (PDB ID: 7BV2) consists of an nsp12-nsp7-nsp8 complex bound to the template-primer RNA and triphosphate form of RTP (11). These data were retrieved from the RCSB Protein Data Bank and subsequently used to obtain the key interactions and contacts between nsp12 and Remdesivir. A total of 56 residues from nsp12 interacted with Remdesivir.

### Molecular docking of EIDD-2801 to the nsp12-nsp7-nsp8 complex bound to the template-primer RNA and triphosphate form of Remdesivir (RTP)

EIDD-2801 is an orally bioavailable NHC prodrug (β-D-N^4^-hydroxycytidine-5’-isopropyl ester) that is similar to Remdesivir, and both drugs work by mimicking ribonucleosides. Therefore, EIDD-2801 was docked into the Remdesivir binding site of the RNA-dependent RNA polymerase of SARS-CoV-2 (15). Molecular docking was performed using the AutoDock Tools to dock the EIDD-2801 into the nsp12 binding cavity, in which it exerts its antiviral action via the introduction of copying errors during viral RNA replication (28, 29). The nsp12-nsp7-nsp8 complex included the RNA and Mg^2+^ catalytic ions. PDB ID: 7BV2 is cocrystallized with the Remdesivir. The same binding grid was selected for the EIDD-2801 docking studies. The protein and ligands were prepared using the MGL tools. Hydrogens and Kollman charges were added to the protein, and hydrogen and Gaisteger charges were added to EIDD-2801 for preparation. The grid was set as X=42, Y=46, and Z=52 with 0.375-Å grid spacing. The protein and ligand were docked, and 100 binding poses were generated for the analysis. The best binding pose that laid well in the binding cavity and showed interactions with the key catalytic residues were selected for further analysis. The interactions between the complex and ligands were identified. A total of 65 residues from nsp12 interacted with EIDD-2801.

### Structural refinement of Remdesivir- and EIDD-2801-bound nsp12-nsp7-nsp8 in complex with template-primer RNA for Rosetta ligand-based interface design

The crystal structure of Remdesivir-bound nsp12-nsp7-nsp8 and the EIDD-2801-bound nsp12-nsp7-nsp8 in complex with template-primer RNA were first optimized using Schrödinger Maestro (Schrödinger Release 2016–4: Maestro, Schrödinger, New York). The parameters for Remdesivir and EIDD-2801 were not available *a priori* for Rosetta-compatible forcefield. Therefore, the appropriate charges were added, and parameters for Remdesivir and EIDD-2801 were generated. The resulting complexes with Remdesivir and EIDD-2801 were used for structural refinement by Rosetta relax, which is the main protocol for all-atom refinement of structures in the Rosetta forcefield (30). Ten models were generated for each design group in this step, namely the Remdesivir-bound nsp12-nsp7-nsp8-template-primer RNA and EIDD-2801-bound nsp12-nsp7-nsp8-template-primer RNA. The full atom-relaxed structures from each group with the lowest energy were used for subsequent ligand-based interface design experiments. The catalytically critical Mg^2+^ ions were retained in all of the refinement and subsequent design steps.

### Ligand-based interface design and binding affinity calculation

Rosetta macromolecular modeling suite was used to perform the ligand-based interface design experiments (31). The nsp12 was redesigned with allowed backbone flexibility using Rosetta’s full-atom scoring function in the Remdesivir- and EIDD-2801-bound nsp12-nsp7-nsp8-template-primer RNA-bound structures. For the Remdesivir-wrapped nsp12-nsp7-nsp8-template-primer RNA-bound structure, 56 residues of nsp12 that are known to interact with Remdesivir were designed with 19 other amino acids, and 65 residues of nsp12 were designed to have intermolecular contacts with EIDD-2801 for the EIDD-2801-bound nsp12-nsp7-nsp8-template-primer RNA-bound structure. During these design experiments, all of the other residues of nsp12, nsp7, and nsp8 were allowed to repack without design. A total of 100,000 designs comprised of Remdesivir- and EIDD-2801-bound complexes were generated. A modified RosettaScripts was used to design the interface residues of nsp12 in complex with Remdesivir and EIDD-2801 that considered small movement of the backbone of nsp12 to avoid steric clashes with ligands upon the introduction of mutations (32). The designs were sampled using Monte-Carlo simulated annealing in the Rosetta all-atom force field. For a comprehensive understanding of the structural and residue-specific changes of nsp12 upon interaction with Remdesivir and EIDD-2801, the Rosetta total score, root mean square deviation (RMSD) from the starting structure, Rosetta Interface delta (representing the binding affinity between the designed nsp12 with Remdesivir and EIDD-2801), and the percentage sequence identity from the starting sequence were obtained.

### Validating the design protocol on Remdesivir-resistant nsp12 mutants of SARS-CoV

To test the accuracy and reliability of our ligand-based Rosetta interface design protocol, we considered a previously reported Remdesivir-resistant mutant of SARS CoV, V557L, in which Val557 is a Remdesivir-interacting residue of nsp12 (15). During this step, the Val557 residue was designed with the 19 other amino acids, but the remaining nsp12 residues were allowed to repack without further design. A total of 5000 designs were generated at 557^th^ position and analyzed to monitor the highly prevalent designed residues with their corresponding scores and rankings.

### Occurrence of mutations in top-ranked affinity-enhanced and affinity-attenuated designs

The type and frequency of designed amino acids in the 56 and 65 nsp12-interacting residues of Remdesivir and EIDD-2801, respectively, from the top-ranked affinity-enhanced and affinity-attenuated designs were obtained and plotted using WebLogo, which graphically represents the multiple sequence alignment of an amino acid profile (33). The residue index is shown in the X-axis of these plots, and the sequence conservation of an amino acid at that position is shown in the Y-axis. The height of symbols within the stack in each plot indicates the relative frequency of a specific amino acid at that position.

### Binding affinity calculation of Remdesivir- and EIDD-2801-bound nsp12 designs using PRODIGY-LIG

The 100,000 Remdesivir- and EIDD-2801-bound nsp12 structures generated using Rosetta ligand-based interface designs were subjected to the calculation of the binding affinities using PRODIGY-LI (PROtein binDIng enerGY prediction - LIGands), which is a structure-based method for the prediction of binding affinity in protein-ligand complexes (34, 35). A 3D model of the design with the bound ligand was supplied, and the binding affinity was computed. From the Rosetta-designed structures, the binding affinity (ΔG) and the contact counts were obtained for analysis. A detailed decomposition of intermolecular contacts was obtained for the top five affinity-enhancing and affinity-attenuating designs for the Remdesivir- and EIDD-2801-bound nsp12 structures.

### Calculation of intermolecular interactions between Remdesivir- and EIDD-2801-bound nsp12 designs in the affinity-enhancing and affinity-attenuating mutants

The intermolecular interactions between top-ranked affinity-enhancing and affinity-attenuating Remdesivir-nsp12 and EIDD-2801-nsp12 designs were obtained using the Arpeggio webserver (36). The total number of interactions obtained represents the sum of the number of van der Waals, proximal, polar contacts, hydrogen bonds, aromatic contacts, hydrophobic contacts, and carbonyl interactions.

### Interaction diagrams between Remdesivir-nsp12 and EIDD-2801-nsp12 designs

The 2D-ligand interaction diagrams showing the various types of intermolecular interactions between Remdesivir-nsp12 and EIDD-2801-nsp12 top-ranked affinity-enhancing and affinity-attenuating designs were obtained using top-ranked affinity-enhancing and affinity-attenuating Remdesivir-nsp12 and EIDD-2801-nsp12 designs were obtained using Schrödinger Maestro (Schrödinger Release 2016–4: Maestro, Schrödinger, New York). A distance up to 6 Å from the ligand molecule was considered while constructing the interaction diagrams.

### Normal mode analysis of RdRp nsp12

The normal mode analysis, deformation energies of each mode, calculation of normalized squared atomic displacements, normalized squared fluctuations, and the correlation matrix of nsp12 were calculated using WEBnm@ server (37, 38).

## ACKNOWLEDGMENTS

The authors are grateful to Dr. Kam Y.J. Zhang (Laboratory for Structural Bioinformatics, RIKEN, Yokohama) for his valuable suggestions and continuous support for improving the manuscript. The authors acknowledge RIKEN ACCC for the Hokusai supercomputing resources.

## AUTHORS’ CONTRIBUTIONS

AKP carried out all the design experiments, data generation, and analysis. RS performed the docking experiment and design file preparation. AKP and TT conceived the study and participated in its design and coordination. AKP, RS, and TT analyzed the data and drafted the manuscript. All authors read and approved the final manuscript.

