## Supplementary Figures for "Rational Design of the Remdesivir Binding Site in the RNA-dependent RNA Polymerase of SARS-CoV-2: Implications for Potential Resistance"

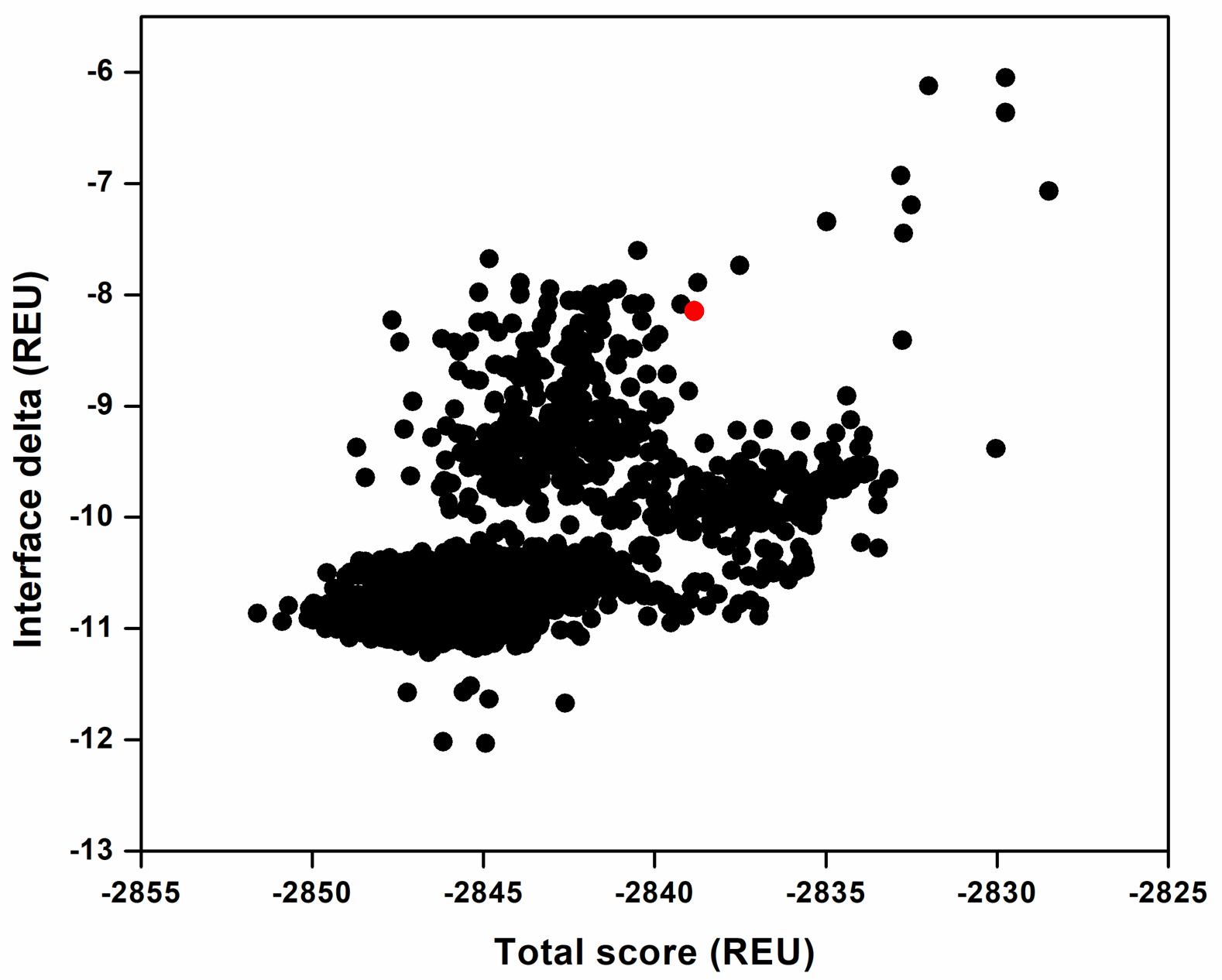

**Fig. S1.** **Rosetta total scores versus Interface deltas of the computed designs where only V557 was designed.** The V557L design was among the low-scored designs shown in red color.

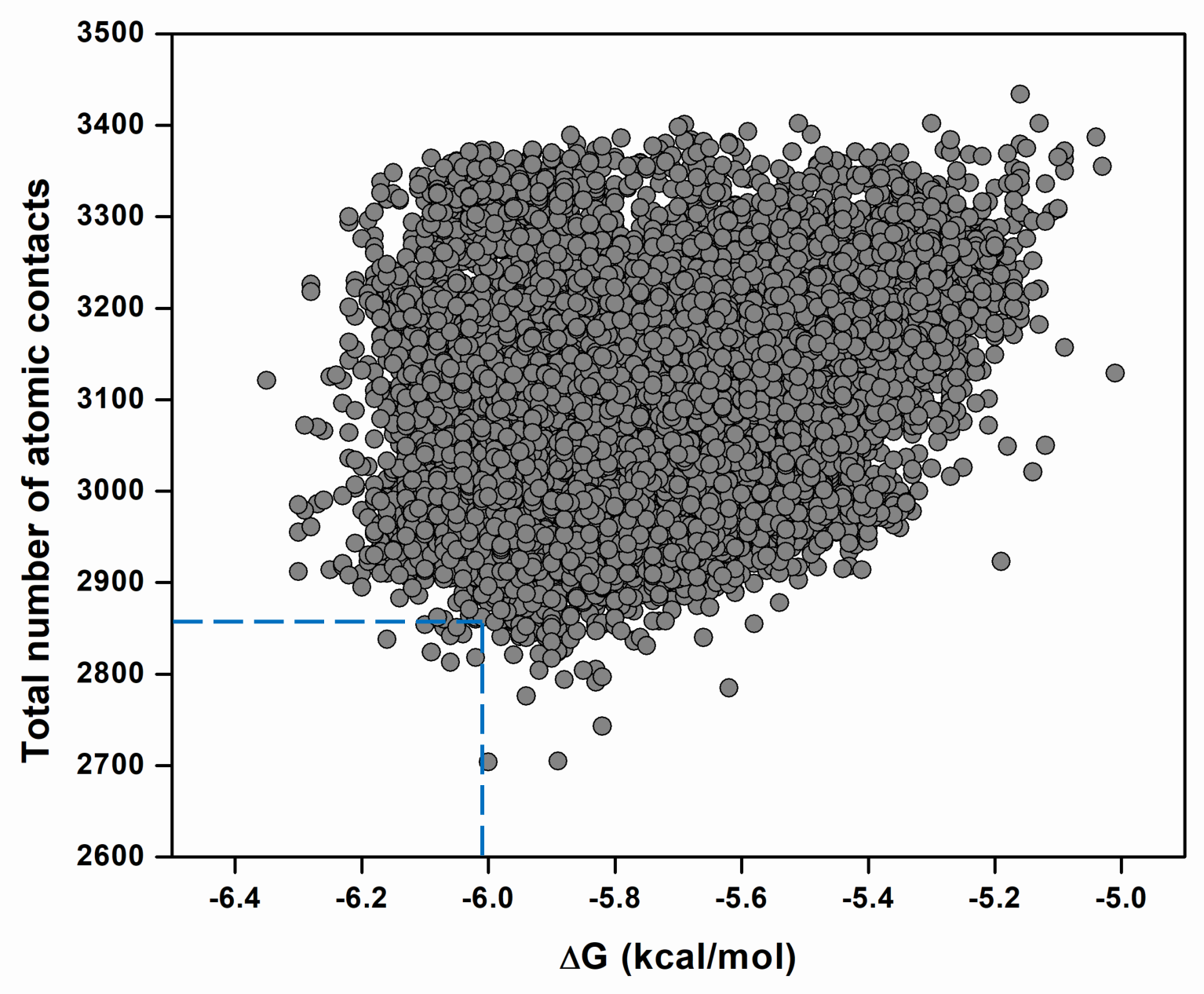

**Fig. S2. PRODIGY-LIG-derived binding affinities and atomic contacts.** The binding affinities between Remdesivir and the nsp12-nsp7-nsp8-RNA complex for the 50,000 generated designs are shown. The contact count represents the summation of (CC+CN+CO+CX+NN+NO+NX+OO+OX+XX) atomic contacts.

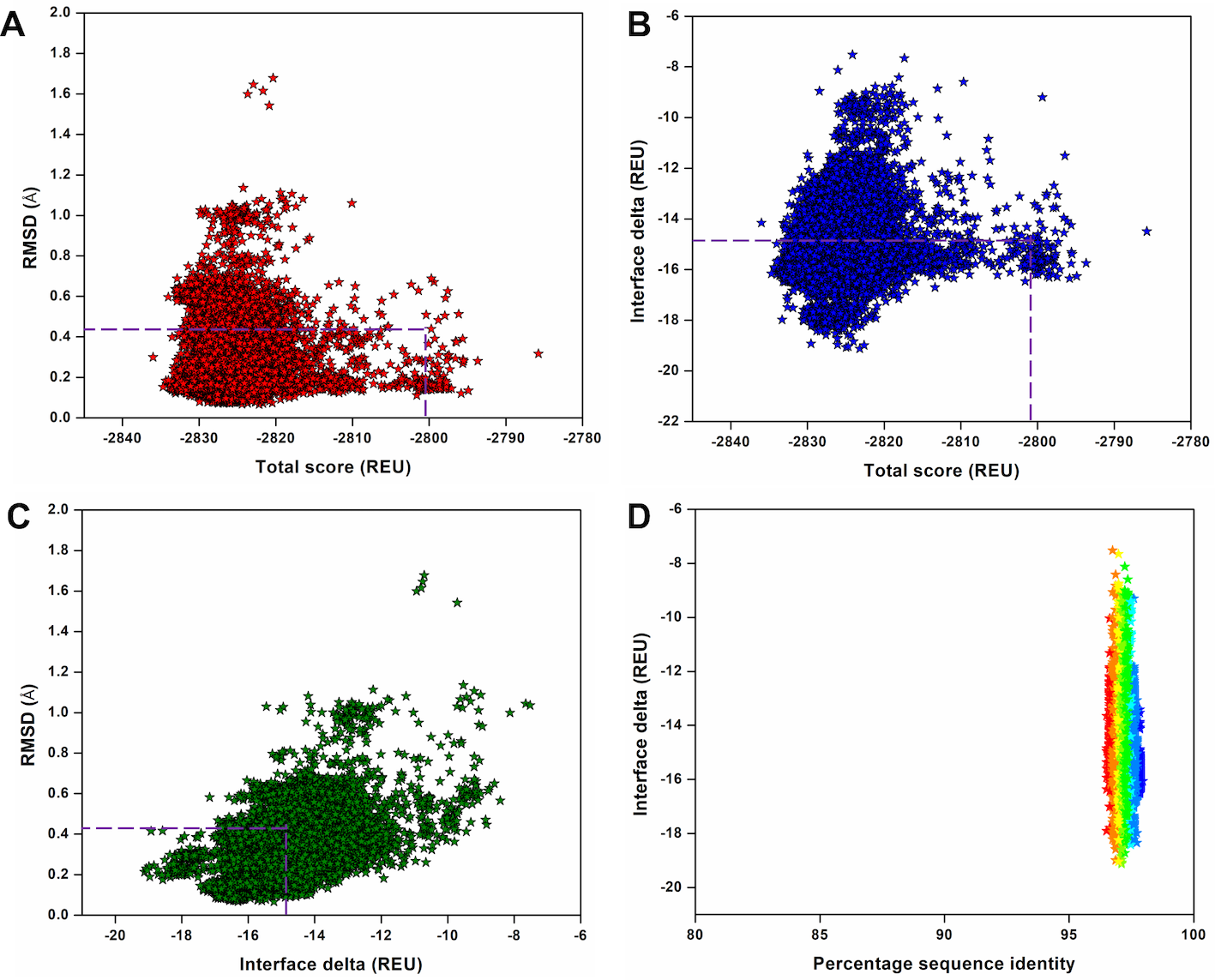

**Fig. S3. Structural and physicochemical parameters from the ligand-based Rosetta interface design experiment.** (A) Rosetta total score versus RMSD of the 50,000 designs of the EIDD-2801-bound nsp12-nsp7-nsp8 complex bound to the template-primer RNA complex obtained from the ligand-based interface design. (B) Rosetta total score versus Interface delta representing the binding affinities of the designs between EIDD-2801 and the nsp12-nsp7-nsp8-RNA complex. (C) Interface delta versus RMSD of the designs, in which only the EIDD-2801-interacting 65 residues were designed, and the remaining residues were repacked. (D) Interface delta versus percentage sequence identity of the EIDD-2801 interacting residues of the nsp12-nsp7-nsp8-RNA complex. In panels (A-C), the dotted boxes denote the control values in which the EIDD-2801-interacting residues were only repacked and not designed.

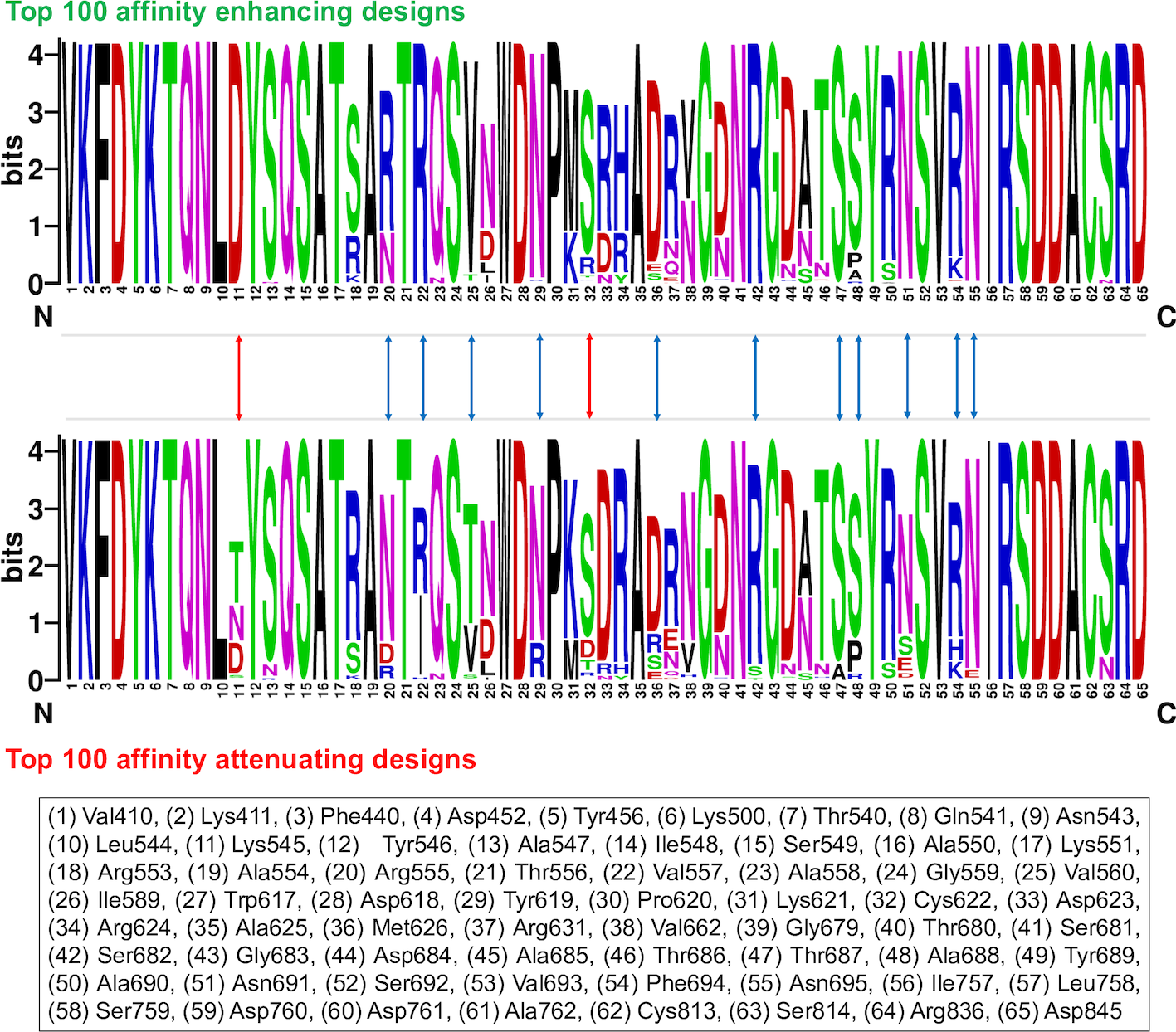

**Fig. S4. Sequence logos showing the frequency of the designed EIDD-2801-interacting residues.** Sequence logos of the 100 top-scored affinity-enhancing versus affinity-attenuating designs generated from the ligand-based interface design of the EIDD-2801-bound nsp12-nsp7-nsp8-RNA complex are shown. The integers on the X-axis represent corresponding native residues, and their identities are shown in the bottom panel. The ‘bits’ represent the overall height of the stack in the Y-axis with the sequence conservation at that position. The blue arrow shows residues that exhibited diverse mutations between the two groups but with a fewer number of amino acids sampled, and the red arrow shows residues that experienced more diverse sequence variations between the two groups with a relatively higher number of sampled amino acids.

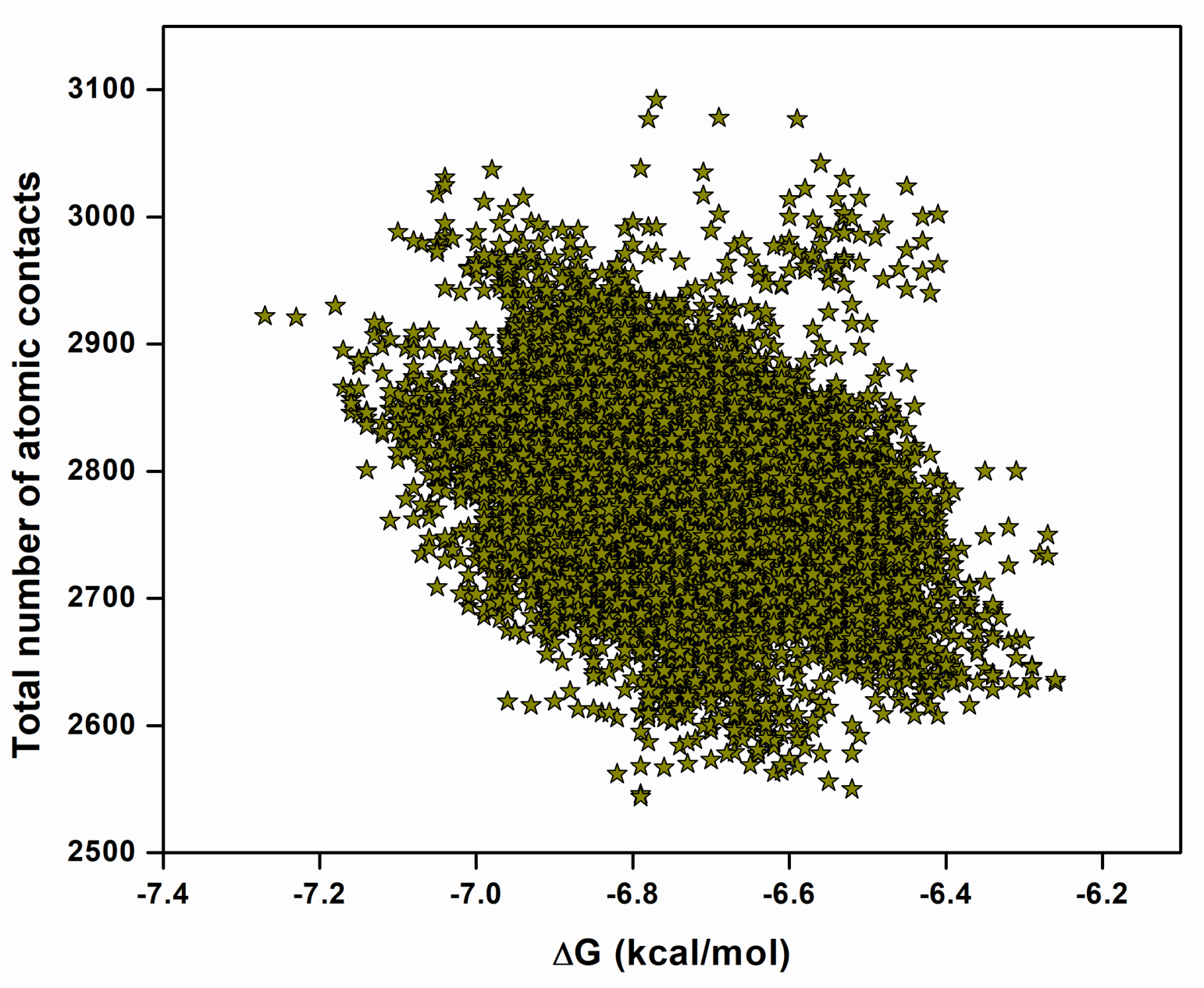

**Fig. S5. PRODIGY-LIG-derived binding affinities and atomic contacts.** The binding affinities between EIDD-2801 and the nsp12-nsp7-nsp8-RNA complex for the 50,000 generated designs are shown. The contact count represents the summation of (CC+CN+CO+CX+NN+NO+NX+OO+OX+XX) atomic contacts.

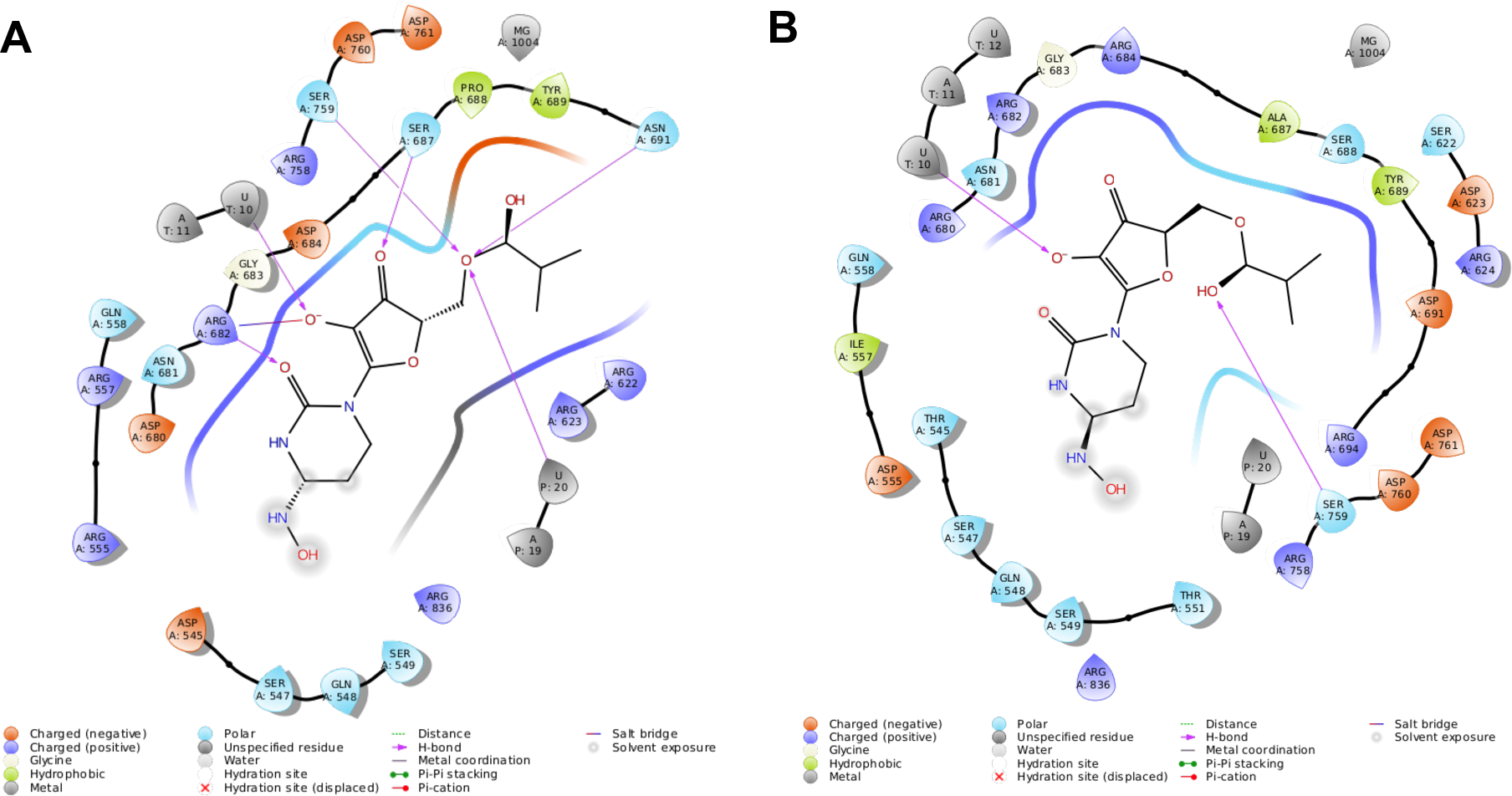

**Fig. S6. Ligand interaction diagrams between EIDD-2801 and the nsp12-nsp7-nsp8-RNA complex for the top-scored affinity-enhancing and affinity-attenuating designs.** 2D ligand interaction diagrams between EIDD-2801 and the nsp12-nsp7-nsp8-RNA complex for the top-scored (A) affinity-enhancing design and (B) affinity-attenuating design are shown within a 6-Å distance. Various types of intermolecular interactions are labeled as legends.

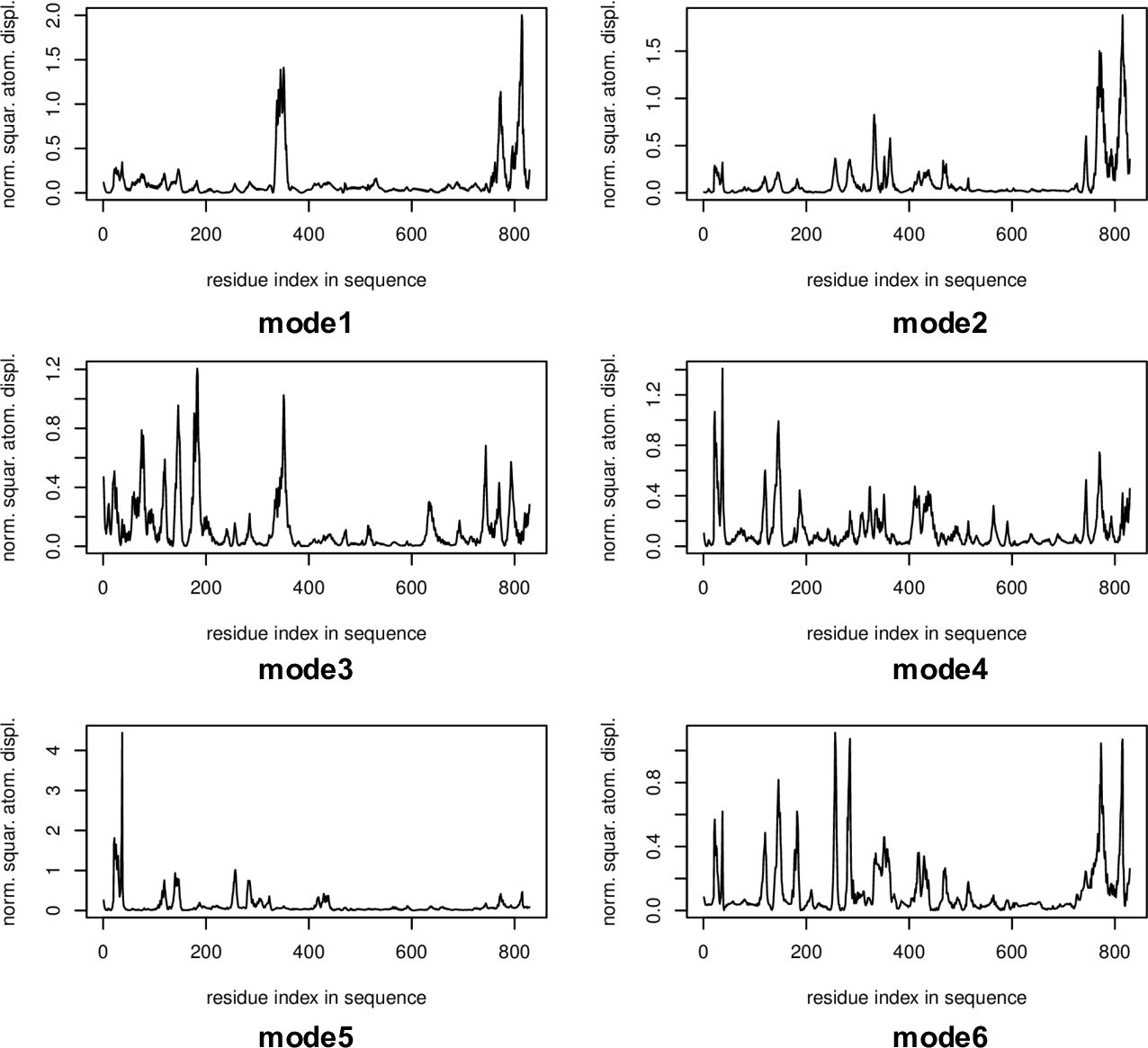

**Fig. S7. Normal mode analysis of RdRp nsp12.** The normal mode analysis and root mean squared atomic displacements of nsp12 for six-modes are shown.
